# Genomic analyses of the extinct Sardinian dhole (*Cynotherium sardous*) reveal its evolutionary history

**DOI:** 10.1101/2021.02.26.432714

**Authors:** Marta Maria Ciucani, Julie Kragmose Jensen, Mikkel-Holger S. Sinding, Oliver Smith, Saverio Bartolini Lucenti, Erika Rosengren, Lorenzo Rook, Caterinella Tuveri, Marisa Area, Enrico Cappellini, Marco Galaverni, Ettore Randi, Chunxue Guojie, Guojie Zhang, Thomas Sicheritz-Pontén, Love Dalén, M. Thomas P. Gilbert, Shyam Gopalakrishnan

**Affiliations:** Section for Evolutionary Genomics, GLOBE Institute, University of Copenhagen, Copenhagen, Denmark; Department of Health Technology, Technical University of Denmark, Kongens Lyngby, Denmark; Smurfit Institute of Genetics, Trinity College Dublin, Dublin 2, Ireland; Micropathology Ltd, University of Warwick Science Park, Coventry, UK; Dipartimento di Scienze della Terra, Paleo[Fab]Lab, Università di Firenze, Via G. La Pira 4, 50121, Firenze, Italy; Sezione di Geologia e Paleontologia, Museo di Storia Naturale, Università degli Studi di Firenze, Via G. La Pira 4, 50121, Firenze, Italy; Department of Archaeology and Ancient History, Lund University, Helgonavägen 3, Box 192, 221 00 Lund, Sweden; Soprintendenza Archeologia, Belle Arti e Paesaggio per le province di Sassari e Nuoro (Ufficio Operativo di Nuoro), Via G. Asproni 8, 08100, Nuoro, Italy; WWF Italy, Science Unit, Rome, Italy; Department of Chemistry and Bioscience, Faculty of Engineering and Science, University of Aalborg, Aalborg, Denmark; BGI-Shenzhen, Shenzhen, 518083, China; Villum Center for Biodiversity Genomics, Section for Ecology and Evolution, Department of Biology, University of Copenhagen, Denmark; State Key Laboratory of Genetic Resources and Evolution, Kunming Institute of Zoology, Chinese Academy of Sciences, Kunming, 650223, China; Center for Excellence in Animal Evolution and Genetics, Chinese Academy of Sciences, 32 Jiaochang Donglu, Kunming 650223, China; Centre of Excellence for Omics-Driven Computational Biodiscovery (COMBio), Faculty of Applied Sciences, AIMST University, Batu 3 1/2, Butik Air Nasi, 08100 Bedong, Kedah, Malaysia; Centre for Palaeogenetics, Svante Arrhenius väg 20C, 10691 Stockholm, Sweden; Department of Bioinformatics and Genetics, Swedish Museum of Natural History, Box 50007, 10405 Stockholm, Sweden; Norwegian University of Science and Technology, University Museum, Trondheim, Norway

## Abstract

The Sardinian dhole (*Cynotherium sardous*)^1^ was an iconic and unique canid species of canid that was endemic of Sardinia and Corsica until it became extinct at the end of the Late Pleistocene^2–5^. Given its peculiar dental morphology, small body size and high level of endemism, several canids have been proposed as possible ancestors of the Sardinian dhole, including the Asian dhole and African hunting dog ancestor ^3,6–9^. Morphometric analyses^3,6,8–12^ have failed to clarify the evolutionary relationship with other canids.

We sequenced the genome of a *ca* 21,100 year old Sardinian dhole in order to understand its genomic history and clarify its phylogenetic position. We found it represents a separate taxon from all other living canids from Eurasia, Africa and North America, and that the Sardinian and Asian dhole lineages diverged *ca* 885 ka. We additionally detected historical gene flow between the Sardinian and Asian dhole lineages, that ended approximately 500-300 ka, when the landbridge between Sardinia and mainland Italy was broken, severing their population connectivity. Our sample showed low genome-wide diversity compared to other extant canids - probably a result of the long-term isolation - that could have contributed to the subsequent extinction of the Sardinian dhole.

## Results & Discussion

We successfully resequenced the genome of a Sardinian dhole (SD) specimen from Corbeddu Cave (Sardinia) (Fig 1A-B) to an average coverage of *ca*. 5x (Table S2). This sample has been radiocarbon dated to ca 21,137 calibrated years before present (Fig 1C) and shows high DNA damage levels (Fig S1A-C).

**Figure 1.**
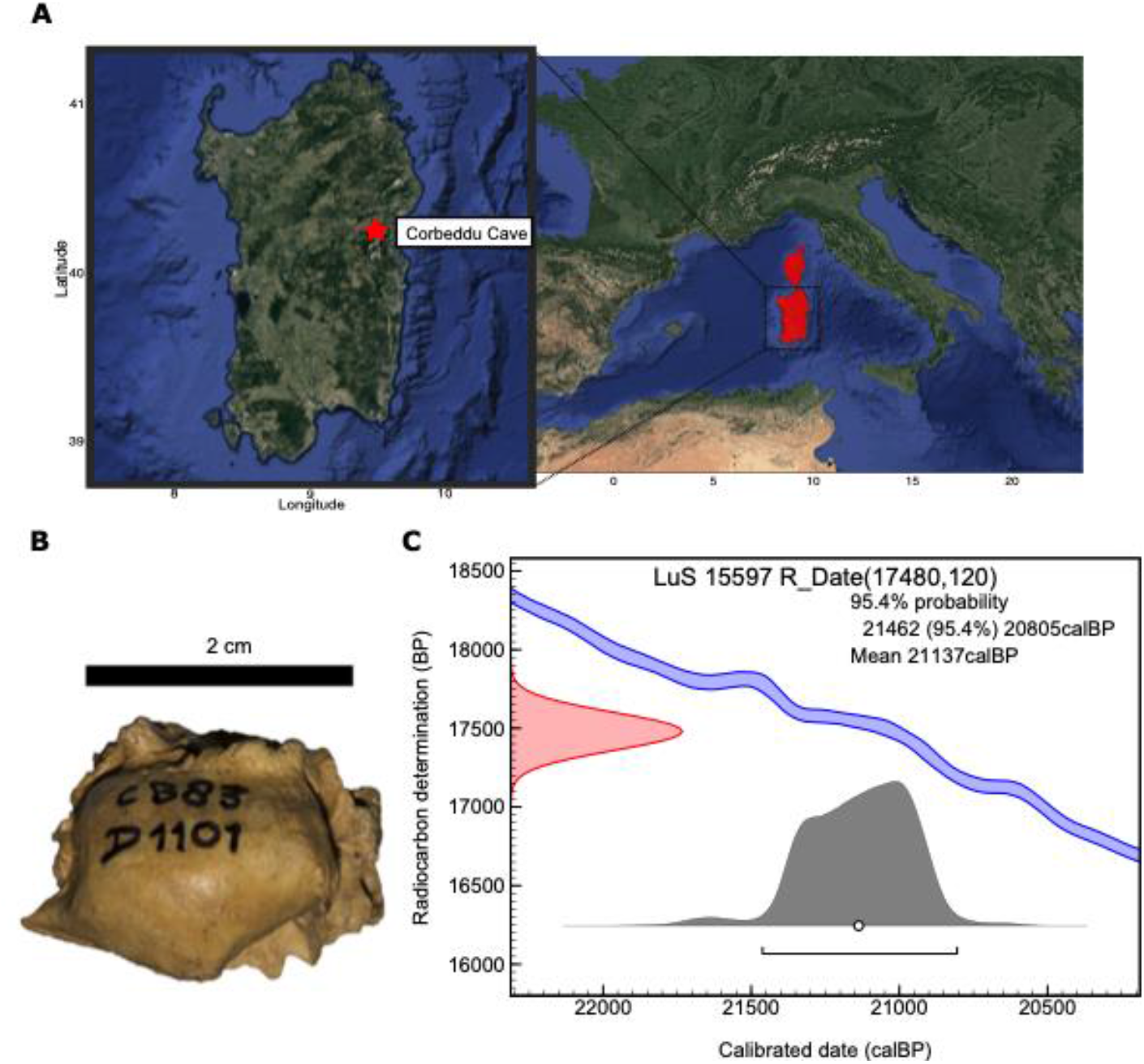
Information related to the sample. A) Sampling location (inset) and hypothetical distribution range of *Cynotherium*. Inset: Exact location of the archaeological site (Corbeddu Cave) in which the Sardinian dhole was excavated. B) Picture of the petrous bone analysed in this study. C) Radiocarbon-dating results of the petrous bone fragment.

Analysis of the reads coverage at each chromosome revealed that the Sardinian dhole specimen was female (Fig S1-E).

To place the Sardinian dhole in an evolutionary context along other canids, we analysed it together with 46 previously published canid genomes (Table S3). The samples used as reference dataset span from 4.4X to 28.2X genome wide coverage and cover the species diversity of the Eurasian, African and American wolf-like canids, including African hunting dogs (*Lycaon pictus*), Asian dholes (*Cuon alpinus*), an Ethiopian wolf (*Canis simensis*), coyotes (*Canis latrans*), African golden wolves (*Canis lupaster*), golden jackals (*Canis aureus*), grey wolves (*Canis lupus*), domestic dogs (*C. l. familiaris*) and an Andean fox (*Lycalopex culpaeus*) as outgroup. To assess the genetic relationships of the extant canids with the Sardinian dhole, we first performed a principal component analysis (PCA) on 46 individuals (excluding the outgroup). Consistent with previous studies ^13^, we found that along PC1 (55.9%) and PC2 (8.45%), the African hunting dogs (AHDs) cluster together and are differentiated from the other canids included in this study. At the same time, the Sardinian dhole holds a distinct placement, whereby, along the first two principal components, it is placed near the two representative modern Asian dholes (Fig 2A). The second principal component separates the genus *Canis* from the dhole samples. Similarly, when we performed PCA excluding the AHDs, the first component placed the Sardinian dhole between the Asian dhole and the *Canis* group (Fig S2-A).

**Figure 2.**
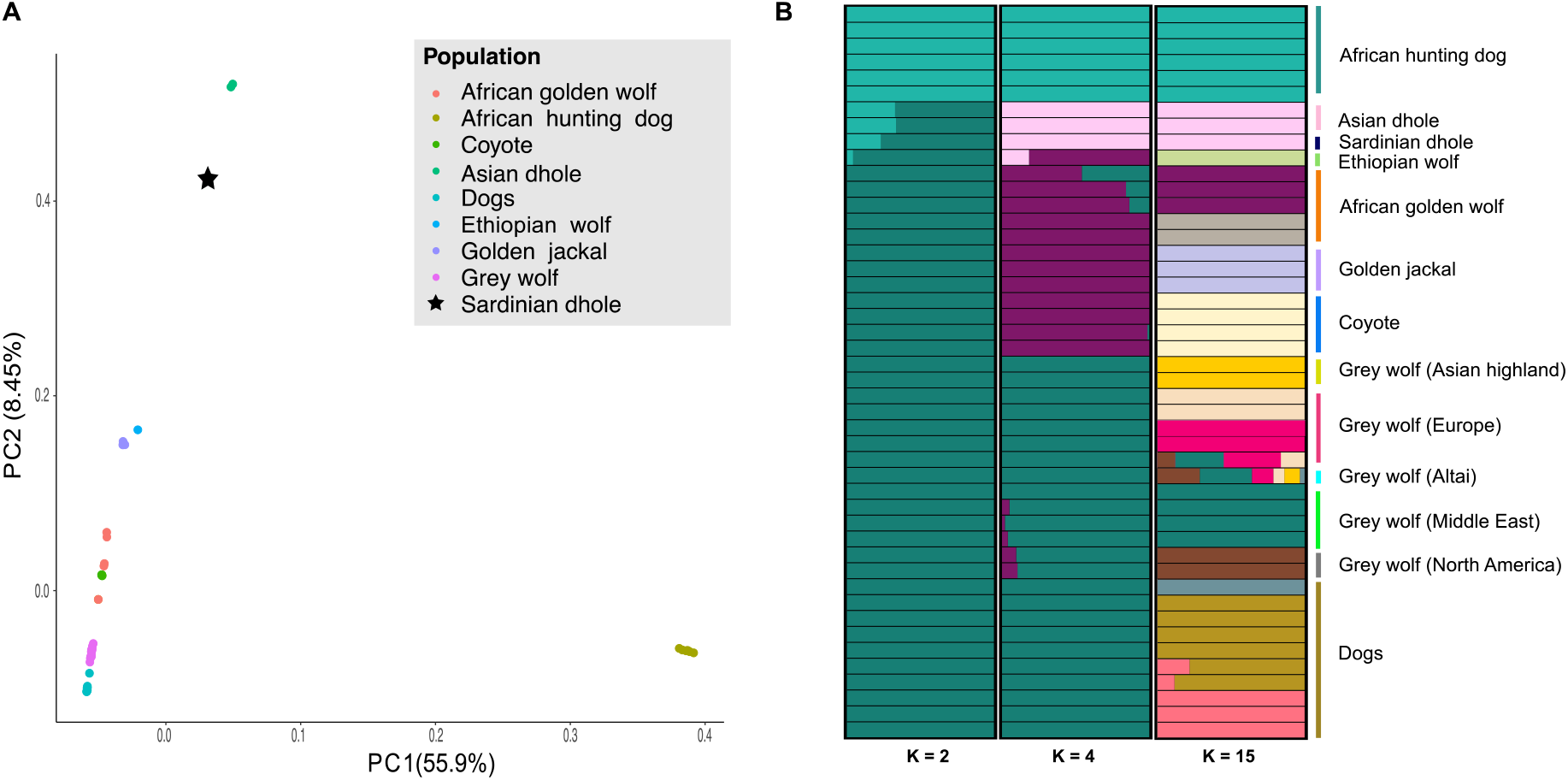
The structure of the canids genomic diversity. A) Principal component analysis of the Sardinian dhole with canid species from Eurasia, Africa and North-America. The black star represents the individual of *Cynotherium sardous* analysed in this study. B) Admixture ancestry component analysis selected for 46 individuals belonging to eight canid species. Only the run with the best likelihood out of 100 runs was selected to be displayed for each K.

To further investigate the relationships between the Sardinian dhole (*Cynotherium sardous*) and other canids, we performed an admixture analysis based on genotype likelihoods for all the samples excluding the Andean fox (Fig. 2B and Fig S2-B). Clearly, when only two ancestral clusters (K=2) were estimated, the most basal canids - AHDs - separated from the *Canis* genus, with the two dhole species being represented as mixtures of these two clusters. Upon increasing the number of estimated ancestry clusters to four (K=4), we observed a division between the AHDs, the dholes, wolves/dogs and the rest of the canids. Specifically, the AHD maintains the same structure while the Sardinian dhole and the Asian dholes are grouped into a cluster of their own. Upon increasing the number of estimated ancestry clusters, the samples fall in clusters according to species and/or populations, with the Sardinian dhole clustering with the two Asian dholes.

To explore the phylogenomic placement of the Sardinian dhole among other canids, and especially understand its relationship with the Asian dhole, we used ASTRAL-III^14^ to estimate the species tree of the canids included in this study by combining 1000 gene trees estimated from randomly chosen 5kb regions across the nuclear genome. The estimated species tree was rooted using the Andean fox as the outgroup (Fig S2-C). In the multispecies coalescent tree estimated by ASTRAL-III, the Sardinian dhole forms a distinct clade, inside the Asian dhole and sister to *Canis*. We subsequently tested the discordance between the species tree and the gene trees to quantify the uncertainty of the branch that split the Sardinian dhole, Asian dhole, *Canis* and the basal canids. The frequencies of the three bipartitions induced by the aforementioned branch (identified as branch 16 in Fig 3A) are shown in Fig 3B, along with similar measures for all the internal branches of the species tree. Two of the three possible bipartitions induced by branch 16 have a frequency greater than 33% - the cutoff previously shown to be required for identifying the true topology^15^. Essentially, although gene trees clustering the Sardinian dhole the Asian dholes are more likely, both topologies, i.e. Sardinian Dhole clustering with Asian dholes or *Canis* are observed in more than 33% of all gene trees, implying that both these topologies could represent the true phylogeny.

**Figure 3.**
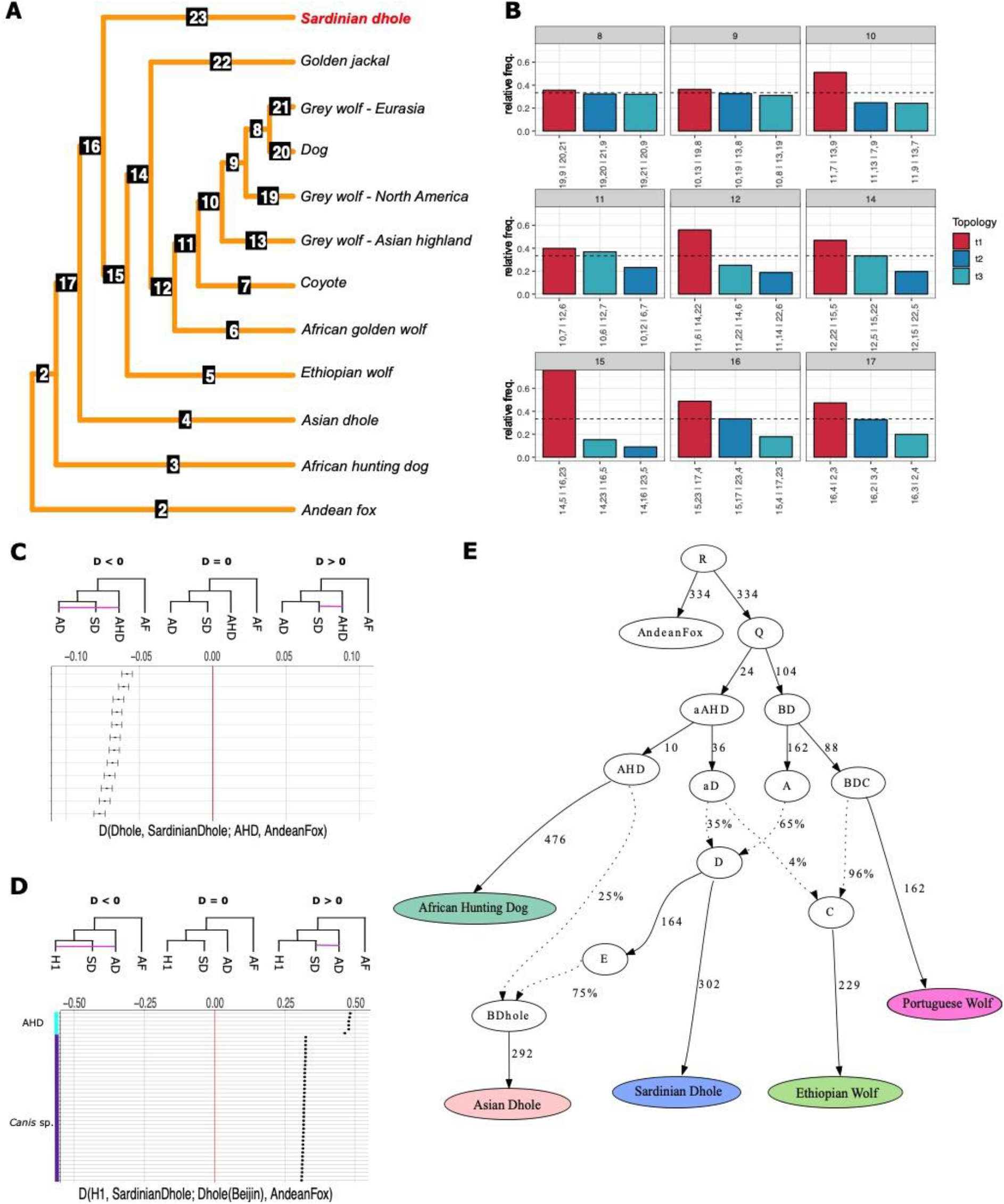
Whole genome phylogeny, gene flow and population ancestry model of the Sardinian dhole. A) Species tree phylogeny generated by Astral-III estimated for 1000 genomic regions. The tree was rooted on the Andean fox (*Lycalopex culpaeus*). Monophyletic clusters were collapsed into the same leaf node. B) DiscoVista relative frequency analysis. Each box title indicates the corresponding branch on the tree in panel A and shows the frequency of three topologies around the focal internal branch of ASTRAL species tree. The first topology represented in red is the main topology followed by the other 2 alternatives in blue. On the y-axes the relative frequency is indicated and the dashed lines represent the ½ threshold. On the x-axis each quartet topology is shown using the neighboring branch labels. C-D) In these panels the gene flow among different canids is shown using ABBA-BABA test in ANGSD. On the y-axes of panel C) different combinations of Asian dholes (AD) and AHD individuals are considered (e.g. from the top to the bottom dhole (Berlin zoo)-AHD (Kenya), dhole (Beijing zoo)-AHD (Kenya) and so on). In the panel C) there is significant gene flow between the Asian dhole (AD) and the African hunting dog (AHD) showing a higher degree of genetic affinity between these two groups compared to the Sardinian dhole (SD) and AHD. In panel D) allele sharing between the AD and SD is higher than when considering AD with any of the other canid lineages in the study. E) Model of the phylogenetic relationships among canids augmented with admixture events. The qpGraph shown here was estimated considering the pairwise D-statistics. Dotted lines represent admixture events and the estimated mixture proportion is shown along them (%). Genetic drift (expressed in drift units per 1,000) is shown along solid lines. This admixture graph represents the best fitting graph (−3 > Z < +3) to model African hunting dog-like ancestry into the Asian and Sardinian dhole and Ethiopian wolf.

The uncertain placement of the Sardinian dhole on the phylogenetic tree led us to investigate historical gene flow events between this species and the other canids. D-statistics, which uses quartets of populations/samples, were used to identify gene flow between canid lineages. We computed the D-statistics using all possible triplets of samples with the Andean fox as the outgroup (Table S4).

Considering the past distribution range of the AHD and dholes’ ancestors, we focused on the topology in which the AHD is the sister clade, with the Sardinian dhole and Asian dhole in the ingroup, i.e. (((Asian dhole, Sardinian dhole), AHD), Andean fox). Our results (D=−0.068, z-score=-19.41, Table S4) suggest significant excess of allele sharing between the Asian dholes and the AHD compared to the allele sharing between the AHD and the Sardinian dhole. Next, we looked for signs of gene flow between the Sardinian dhole and other species of *Canis*. When testing the following three different combinations: 1. (((*Canis*, Golden Jackal), Sardinian dhole), Andean fox), 2. (((*Canis*, Ethiopian wolf), Sardinian dhole), Andean fox) and 3. (((*Canis*, African golden wolf), Sardinian dhole), Andean fox) the D-statistics suggest higher allele sharing between *Canis lupus* and the Sardinian dhole (Fig S3). This result could arise from two scenarios; 1. There was indeed gene flow between the grey wolf and the Sardinian dhole, or 2. The trio of related species - Ethiopian wolf, African golden wolf and Golden jackal - share ancestry with a species not represented in our study, that falls outside the Sardinian dhole in the phylogeny.

Assuming that the ancestor of the *Cynotherium* arrived in Sardinia-Corsica through a terrestrial connection between the islands and mainland Europe, this sets the possible colonization of Sardinia-Corsica at *ca* 5 Ma, during the Messinian salinity crisis ^16^, or around 3 Ma - close to the Plio-Pleistocene boundary ^12,17^ However, since then, there is no proof of a landbridge that would allow movements of species between the two islands and the continent. Nevertheless, Pleistocene mammals of Sardinia are currently divided into two major faunal complexes, an older one (*Nesogoral* Faunal Complex) and a younger one (*Tyrrhenicola* Faunal Complex) divided by the end of the Early Pleistocene ^18,19^. Given this complex scenario, a number of earlier studies considered the Sardinian dhole as a subgenus of *Xenocyon, Cuon*, or a derived form of *Canis* ^3,10–12^ that might have reached Sardinia and Corsica by sweepstake or passive dispersal at the transition between Early and Middle Pleistocene ^4–20^. This phenomenon is especially feasible during periods of fluctuation of the sea level and have been known to contribute to the faunal turnover in Sardinia^4,18,20^ Therefore, to further investigate the demographic history of dholes, we used admixture graphs to test whether the diffuse ancestries can be explained by AHD-like admixture in Asian dhole or *Canis*-like admixture in Sardinian dhole. First, we estimated the proportion of ancestry derived from the ancestor of the Asian dhole into the Sardinian dhole using AHD, dholes, Ethiopian wolf and Andean fox as outgroup. We find that the node representing the ancestral population of the Asian and Sardinian dholes derives 60% ancestry from the node that diverged from the ancestral AHD population in the past (Fig 3E). A second admixture event brings 25% of AHD ancestry into only the ancestor of the Asian dholes. Subsequently, when including the Portuguese wolf in the previous graph we found that it was necessary to model one more admixture event (see Fig S4) between the ancestors of the Ethiopian wolf and the dholes. The Ethiopian wolf is best modelled as a mixture of 4% from an ancestral population related to the dholes and 96% from the *Canis* lineage. This last admixture event confirms the results obtained in admixture using four ancestry components, (Fig 2B) in which the Ethiopian wolf shares a proportion of its ancestry with the dholes.

Given the geological history of Sardinia and Corsica, along with cycles of long-term isolation and past colonization, we explored the effects of isolation on genetic diversity in the Sardinian dhole using estimates of genome-wide heterozygosity and effective population size. We thus first inferred the heterozygosity in sliding windows for all the representative canid species in our dataset and we found that the Sardinian dhole shows remarkably low levels of heterozygosity across the whole genome, comparable to other isolated canids with small population sizes, such as the Ethiopian wolf and the AHD (Fig. 4A).

**Figure 4.**
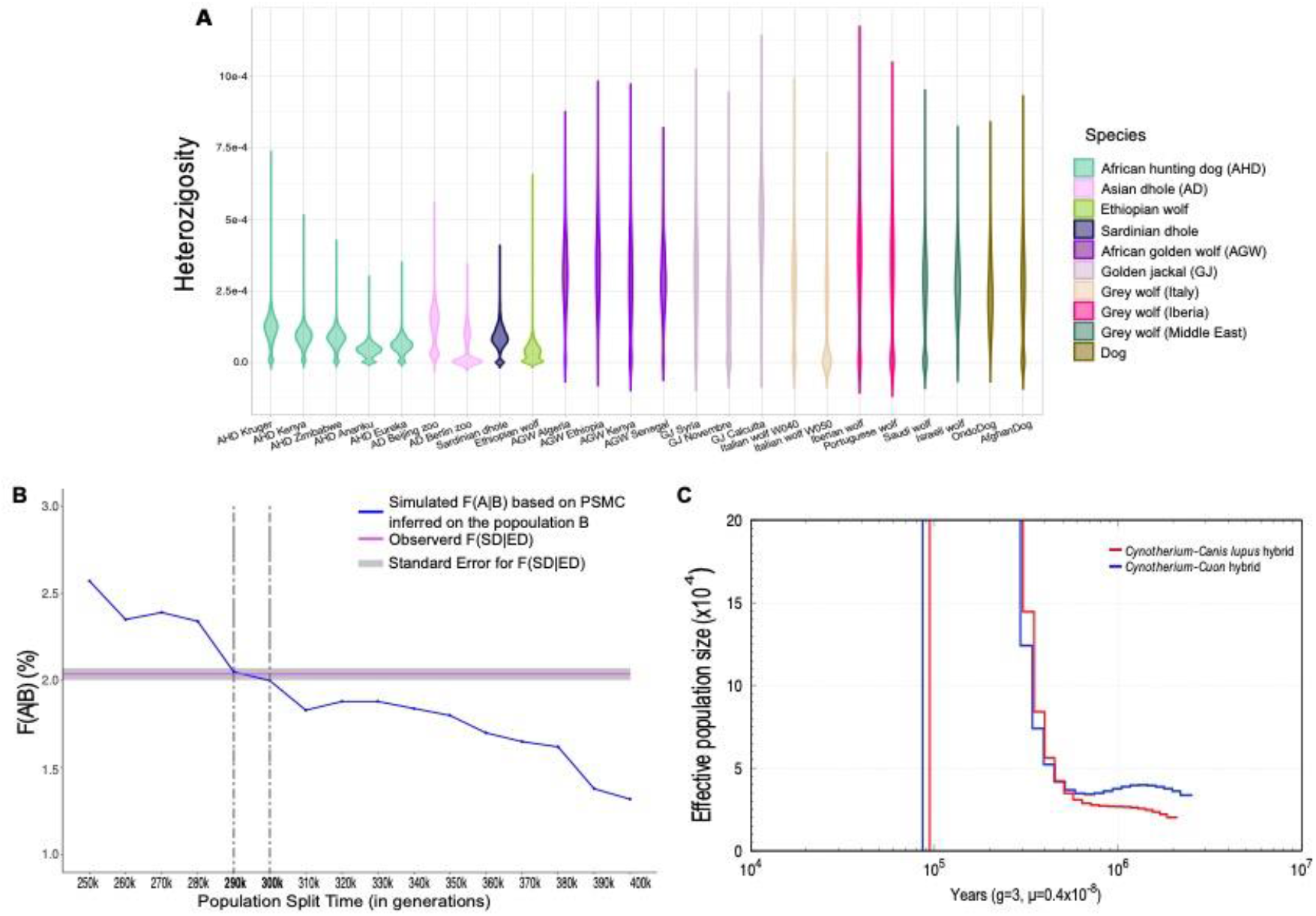
Population dynamics, split time and end of gene flow. A) Violin plot representing heterozygosity in sliding windows across the species of canids selected in this study. B) Estimate of the population split time between the Sardinian dhole and Asian dhole populations. F(A|B), represented on the y-axes, is the probability to observe a derived allele in the population A (SD) when B (ED) is heterozygous at the same site. The numbers on the x-axis represent the population split time expressed in generations. C) hPSMC plot based on the artificial hybrid genomes constructed using different species.

The other two dholes show a strongly bimodal distribution in their heterozygosity distribution, with regions of high and very low heterozygosity, probably as a result of their recent demographic histories and captivity breeding. Among the different canid species that went through bottlenecks or population reductions ^21–26^, the Sardinian dhole shows decreased genetic diversity across the entire genome, comparable to the AHDs from Zimbabwe, Kenya and South Africa (Fig. 4A) that went through a stable and long-term population decline ^27,28^. This result represents a clear picture of the past population dynamics of the Sardinian dhole and how the long-term isolation has shaped its genome. The low levels of heterozygosity in the Sardinian dhole across all regions of its genome strongly suggest a long period of time with low effective population size, exhibiting population dynamics similar to that of the mountain gorilla ^29^.

We then calculated the divergence time between the Sardinian dhole and the contemporary Asian dhole using the statistic F(A|B), which estimates the probability of an individual A (Sardinian dhole) carrying the derived allele at sites that are heterozygous in individual B (Asian dhole). The assumption behind this approach is that when two populations start to diverge, they will also accumulate mutations that - due to isolation - will not be shared with other populations. We estimated that the Sardinian dhole carried the derived allele at ~2% of sites that were heterozygous in the Asian dhole. The proportion of derived alleles was then used as a summary statistic to estimate the divergence time of the Sardinian dhole and the Asian dhole from simulations of multiple divergence times using the population history of the Asian dhole inferred from PSMC. The simulations were calibrated using the wolf mutation rate (μ = 4 × 10^−9^ /bp/generation) ^30^ and a generation time (g) of 3 years ^30,31^. Using F(A|B), we conclude that the Sardinian dhole and the Asian dhole diverged *ca* 885 ka (range 870 ka and 900 ka) (Fig. 4B).

Lastly, explored the timing of gene flow between the Sardinian/Asian dhole and Sardinian dhole/Eurasian wolves and dogs by performing a pseudodiploid demographic analysis using hPSMC^32^. The results suggest that the gene flow between the Sardinian dhole and the ancestor of the modern Asian dhole ended between 510 ka and 310 ka, while gene flow between the Sardinian dhole and the Eurasian wolf-like ancestry ended between 510 ka and 360 ka. This shows that gene flow between the Sardinian dhole and the other two lineages might have ended around the same time, between 560 ka and 310 ka. We found that the demographic histories of the two pseudo-hybrids (Fig. 4C) are almost identical, indicating that gene flow ceased after the Middle Pleistocene, a timing that is also consistent with a number of publications in which different faunal complexes that populated the island were taken into consideration ^18,20,33^. Further, we determined that estimates using Sardinian dhole/AHD and Asian dhole/AHD hybrids gave an interval between 1.05 Ma and 300 ka for the cessation of gene flow between AHD and the two dhole lineages (Fig S5). Wide intervals, such as those reported here, could be the result of very low levels of post-divergence gene flow between two species ^32^. Furthermore, the last fossil of *Lycaon* found in Europe is dated to 0.83 Ma, implying that this genus went locally extinct in Europe towards the end of the Early Pleistocene ^34^ Combining archaeological evidence and hPSMC analysis, we hypothesize that the divergence time between *Lycaon* and the lineage that gave rise to the Sardinian dhole and to the modern form of *Cuon* in Europe can be narrowed down to between 1.05 Ma and *ca* 0.83 Ma. Admixture events between the ancestors of the AHD and the Asian dhole lineages could have happened in other locations in the past when the two species were not confined to their present-day ranges. For instance, the first fossil of a *Lycaon* dating back to the Middle Pleistocene was found in Israel, suggesting that the ecological boundaries that separate these two species were not in place yet ^35^. This could explain the 25% ancestry derived from the AHD lineage observed only in the Asian dhole (Fig 3E), which probably occurred after the Sardinian dhole lineage became isolated in Sardinia and Corsica.

In conclusion, by generating the first whole-genome of a Sardinian dhole, we clarified the phylogenomic placement of this species among the genomic diversity of living canids from Eurasia, Africa and North-America. We detected historical gene flow between the Sardinian dhole and Asian dhole lineages, and evidence of past admixture from the ancestor of the AHD into the Asian dhole lineage. We found that the Sardinian dhole lineage diverged from the Asian dholes around *ca* 885 ka, followed by post-divergence gene flow ceasing later, between 560 and 310 ka, meaning that Sardinia could have been colonized several times during the Pleistocene. However, from these results it can be inferred that, probably sometime during the Middle Pleistocene, the physical barrier created by the sea separating Sardinia from the continent effectively ended further canid migrations. We also found that our sample shows an overall reduced genome wide diversity that - together with the long-term isolation on the island - could have contributed to its extinction. In this complex and delicate situation, the hypothesis that humans played a crucial role in the extinction of the last Pleistocene mammalian fauna cannot be excluded ^36,37^ However, the direct or indirect anthropogenic pressure exerted on the Sardinian dhole population could have been a concause - together with the long-term isolation, loss of genetic diversity and reduction of Ne - leading to the extinction of this species. The results presented here highlight also the importance of sequencing ancient genomes from island ecosystems to improve our knowledge of the past evolution, colonization and migration events of extinct species.

## Methods

### Samples Information

A *Cynotherium sardous* petrous bone (CB83-D1101) from Corbeddu cave was selected at the collection of the National Archaeological Museum of Nuoro. Corbeddu cave is an archeo-paleontological site located in the Lanaittu Valley (Oliena, Nuoro Province, Northeastern Sardinia, 40.254921°, 9.485078°) famous for its lithic and osteological evidence of *H. sapiens* ^38,39^, disputedly referred to the Late Pleistocene or Early Holocene ^40,41^. *Cynotherium sardous* remains, among which a nearly-complete skeleton, come mainly from the upper level of layer 3 of hall II ^3,42^ Such specimens represent the last occurrence of the species, recently re-dated by Palombo and colleagues ^41^ to 12,945 ± 75 years.

### Data generation

The sample was processed under strict clean laboratory conditions at the GLOBE Institute, University of Copenhagen. Small petrous bone chunks of around 350 mg, were divided into 3 eppendorf tubes - A, B and C (*ca* 120 mg each) - and washed with diluted bleach, ethanol and ddH2O, following Boessenkool et al. ^43^.

The subsampled material was extracted following Dabney et al. (2013), purified using modified PB buffer ^44^ and eluted using 2 washes in 22 *μ*l buffer EB (EB) - with 10 minutes of incubation time at 37°C. The concentration of each extracts was checked on Qubit (ng/*μ*l) and on Agilent 2100 Bioanalyzer High-Sensitivity DNA chip (Agilent Technologies) for molar concentration and fragment size. Sequence data was generated using both BGISeq and Illumina sequencing platforms. Specifically, five Illumina libraries were built using 16 *μ*l of DNA from each extract (A, B, C) in a final reaction volume of 40 *μ*l following Carøe et al.^45^ and amplified using PfuTurbo Cx HotStart DNA Polymerase (Agilent Technologies) and Phusion^®^ High-Fidelity PCR Master Mix with HF buffer (New England Biolabs Inc) (see Table S1 for further details). From the extract C two aliquots of 16 *μ*l were used to build BGIseq compatible libraries following Mak et al. ^46^ and Carøe et al ^45^ and amplified with Phusion^®^ High-Fidelity PCR Master Mix with HF buffer (New England Biolabs Inc). The appropriate number of cycles were determined using Mx3005 qPCR (Agilent Technologies) in which 1 *μ*l of SYBRgreen fluorescent dye (Invitrogen, Carlsbad, CA, USA) was loaded in 20 *μ*l indexing reaction volume using also 1 *μ*l of template, 0.2 mM dNTPs (Invitrogen), 0.04 U/μl AmpliTaq Gold DNA polymerase (Applied Biosystems, Foster City, CA, USA), 2.5 mM MgCl2 (Applied Biosystems), 1X GeneAmp^®^ 10X PCR Buffer II (Applied Biosystems), 0.2 μM forward and reverse primers mixture ^46^, and 16.68 μl AccuGene molecular biology water (Lonza). qPCR cycling conditions were 95°C for 10 minutes, followed by 40 cycles of 95°C for 30 seconds, 60°C for 30 seconds, and 72°C for 45 seconds. Library index amplifications were performed in 50 μl PCR reactions that contained 14 μl of purified library, 0.1 μM of each forward (BGI 2.0) and custom made reverse primers ^46^, 2x Phusion^®^ High-Fidelity PCR Master Mix with HF buffer and 8.6 μl AccuGene molecular biology water (Lonza, Basel, CH). PCR cycling conditions were: initial denaturation at 98°C for 45 seconds followed by 18 to 20 cycles of 98°C for 20 seconds, 60°C for 30 seconds, and 72°C for 20 seconds, and a final elongation step at 72°C for 5 minutes. Amplified libraries were then purified using 1.5x ratio of SPRI beads to remove adapter dimers and eluted in 50 μl of EB (Qiagen) after an incubation for 10 minutes at 37°C. Two indexed BGI libraries with the same index combination were pooled together before the beads purification and sequenced on 3 lanes BGIseq-500 using 100 bp single end sequencing reactions. Five Illumina libraries with different combinations of indexes were pooled together in equimolar concentration and sequenced on using 150 bp paired end chemistry on 5 lanes of the Hiseq × platform at SciLifeLab Data Centre, Sweden.

### Radiocarbon Dating

Fragments of the petrous bone of the Sardinian dhole were submitted for AMS ^14^C dating at the Radiocarbon Dating Laboratory at the Department of Geology, Lund University, Sweden. The obtained radiocarbon age, 17,480±120 BP (Lab code LuS-15597), was calibrated based on the INTCAL20 calibration data set ^47^ using the OxCal v4.4.2 software ^48^.

### Dataset

The dataset used in this study is represented by 47 representative canid genomes (Table S3), 46 of which were previously published ^13,27,28,31,49–57^. The genomes considered for this study were chosen to represent the genetic diversity of 9 different species (with the dog considered as a different species from the wolf, Table S3) from Africa, Eurasia, and North America.

### Quality control and alignment

Short reads obtained from BGI and Illumina sequencing platforms were processed using PALEOMIX v1.2.13^58^ pipeline. The same pipeline was run for each sample in the study. In the first step, the adapters were trimmed with AdapterRemoval2^59^ with default settings and the alignments were performed against the wolf reference genome^60^ and the dog reference genome (CanFam3.1)^61^ using BWA v0.7.12 *backtrack* algorithm^62^ with minimum base mapping quality set to 0 to ensure that all the reads were retained in this process. Mapping quality and base quality filters will be applied in the later steps of the analysis. PCR duplicates were filtered out using Picard *MarkDuplicates* v2.9.1^63^ and in the last step GATK v4.1.0.0^64,65^ was used to perform the indel realignment step with no external indel database. Post mortem DNA damage profiles and nucleotide misincorporation patterns (sub C>T and G>A) were computed using mapDamage2.0^66^.

### Sex determination

No Y-chromosome reference sequence is available for CanFam3.1 therefore to infer the biological sex of the Sardinian dhole we used the statistics generated from mapping to the dog reference genome. Rstudio v1.2.5 0 0 3 ^67,68^ and ggplot2 ^69^ were used to visualize the depth of coverage for all the chromosomes of the Sardinian dhole. We then identified as female the individual if the depth of coverage for the × chromosome was similar or exceeded the mean coverage of all the autosomal chromosomes.

### Genotype likelihoods

The samples analysed in the present study span from 4.4X to 28.2X coverage, including in this range also the ancient Sardinian dhole. In order to avoid biases, when possible, we use genotype likelihood over genotype calling. SAMtools model (−GL 1) in ANGSD ^70^ was used to estimate the genotype likelihood at variant sites for all the scaffolds above 1 MB. Bases with base quality lower than 20 and reads with mapping quality lower than 20 were discarded (-minQ 20 -minmapq 20). We retained sites with the default minimum depth for a minimum of 95% of the individuals in our dataset (-minInd 44) and the following parameters: - remove_bads 1 -baq 1 -C 50 -uniqueOnly 1. The resulting output, generated in beagle format, was then used for the PCA and Admixture analysis.

### Principal components analysis

To explore the genetic affinities in our data we performed the principal component analysis (PCA) using PCAngsd v0.98 ^71^ on the 46 individuals’ genotype likelihood panels obtained using ANGSD. A covariance matrix was created and we then used Rstudio to calculate the eigenvectors and eigenvalues on the covariance matrix file and used ggplot2 to plot the PCA.

### Admixture

The genotype likelihood file generated with ANGSD was used as input for NGSadmix v32^72^ to generate the ancestry cluster and proportion of admixture for 46 samples in the dataset using 3008607 SNPs. The outgroup, the Andean fox, was excluded from this analysis and we retained sites with the default minimum depth for a minimum of 95% of the individuals in our dataset (-minInd 44). The structure in the dataset was computed using 2 to 15 clusters (K) and for every cluster the analysis was repeated 100 times to ensure convergence to the global maximum. For each K the replicates with the best likelihood scores were chosen and used as input file list (option -i) in pong ^73^ to visualize the ancestry clusters. The options -n and -l were used to assign the samples’ order and a color for each cluster respectively.

### Heterozygosity in sliding windows

The heterozygosity per sample was estimated using ANGSD, by calculating the per sample folded site frequency spectrum (SFS). We generated a saf.idx file based on individual genotype likelihoods using GATK (-GL 2) from the scaffolds larger than 1Mb (704 scaffolds) for each bam file (doSaf 1 -fold 1) and we excluded transitions (rmtrans 1) and reads with quality score and bases with mapping quality lower than 20 (-minQ 20 -minmapq 20). Since we chose the option -fold 1 to estimate the folded SFS, the wolf reference genome was used both as reference and as ancestral (- ref and -anc options). The repeat regions were masked using a repeat mask of the wolf reference genome ^60^

The scaffolds that were longer than 1 Mb were partitioned into overlapping windows of size 1 Mb with a step size of 500 kb using the *bedtools windows* tool. Windows shorter than 1Mb at the end of the scaffolds were discarded. The SFS for each window was estimated using the realSFS utility tool provided in ANGSD and subsequently the ratio heterozygous sites/total sites was calculated to provide the final heterozygosity per window. R studio ^67^ and ggplot2 and dplyr ^74^ were used to visualize the heterozygosity level at each window and to create a violin plot for a subset of samples.

### Heterozygosity in sliding windows - reference bias and downsampling

To ensure the robustness of our results to the choice of reference genome, we performed the heterozygosity in sliding windows using the same samples mapped to the dog reference genome. Similarly, in order to ensure that the different depths of coverage across the different samples did not lead to significant differences in the heterozygosity estimates, we downsampled 2 individuals, one Italian wolf (ItalianWolf_W050) and one Syrian wolf, of 50% of their original coverage using SAMtools *view* and the option -s 0.5. Heterozygosity in sliding windows was computed on the subsampled data following the same procedure as for the full data. The newly generated bam files were then sorted and indexed using SAMtools *sort* and *index* and went through the same steps used for the original bam files.

### Nuclear genome phylogeny

All the individuals in the dataset, including the outgroup (Andean fox) were used to construct the nuclear genome phylogeny. First, ANGSD was used to generate a consensus sequence for each genome in our dataset using the wolf genome as reference. Each base was sampled based on the consensus base (-dofasta 2). Bases with base quality lower than 20 and reads with mapping quality lower than 20 were discarded (-minQ 20 -minmapq 20). The minimum coverage for each individual was set to 3x (-setminDepthInd 3) and the following additional filters were used: -doCounts 1 -remove_bads 1 -uniqueOnly 1 -baq 1 -C 50. We then selected 1000 random regions, each 5000bp long, from the wolf reference genome using *bedtools random* ^75^ with the following parameters: -l 5000 -n 1000.

Samtools ^76^ was used to generate a fasta file for each region using the consensus sequence generated by ANGSD. For each region, the consensus sequences across all samples were combined into a single multi-sequence fasta file. Subsequently, for each region, RAxML-ng ^77^ was used to reconstruct the phylogeny using the evolutionary model GTR+G. The region-wise gene trees were concatenated together and a species tree was estimated using Astral-III ^14^ with the default parameters, retaining all the branches in the gene trees, i.e. the different individuals of the same species were not collapsed into a single group. Interactive Tree Of Life (iTOL) v4 ^78^ online tool was used to visualise the species tree estimated by Astral-III. Further, we used DiscoVista ^79^ to visualize the discordance between the 1000 gene trees and the species tree generated with ASTRAL-III. For this step, the samples belonging to the same species were then collapsed together using an annotation file (option -a) and the Andean fox was specified as outgroup to root the tree by using the option -g. The option -o was used to create an output folder with the resulting DiscoVista tree and the plot showing the relative frequency analysis which were used to evaluate the three topologies for each internal branch of the tree.

### Gene flow between the Sardinian dhole, AHD and Asian dholes

#### D-statistics (ABBA-BABA)

The program ANGSD was used to investigate the presence and extent of gene flow between the Sardinian dhole and the other canids included in this study. We restricted our analyses to only the scaffolds over 1 MB. To compute the D-statistics using ANGSD, sites with base quality and mapping quality lower than 20 (-minQ 20 -minMapQ 20) were discarded, and at each site a single allele was randomly sampled (-doAbbababa 1). Transitions were also removed (-rmTrans 1) from the analysis to avoid biases introduced by ancient DNA damage and the following options were used: -doCounts 1 -useLast 1 -blockSize 1000000. We tested all triplets of samples, using the Andean fox as the outgroup. The subset of triplets representing the correct tree topology, as estimated by ASTRAL-III and DiscoVista, were considered for testing gene flow hypothesis between the Sardinian dhole and the other species. Using a similar approach, we also investigated gene flow between Asian dholes and AHDs. Finally, D-statistics with a Z-score between 3 and −3 were not considered significant.

#### Calling of polymorphism and filtering

We used GATK v3.4.0 with the option -T*Haplotype caller* ^80^ tool to call variants for all the samples in our dataset, using the wolf genome as reference (option -R). The option -L was used to specify the regions of our interest (scaffold above 1 Mb) and sites with base quality and mapping quality lower than 20 (-mmq 20 -mbq 20) were discarded. The resulting *vcf* files were then compressed and validated using VCFtools v0.1.8 ^81^ option *VCF-validator*. VCFtools was subsequently used for further filtering the VCF files by excluding indels, sites with minimum depth below 5 (-minDP 5). All the sites with more than 90% of missing genotypes (-max-missing-count) over all individuals were excluded and the option -plink was used to generate the plink files in MAP and PED format. The plink files were then merged using Plink v1.9 ^82,83^ and used to generate the files BED BIM and FAM. The following options were applied: -merge-list, -allow-extra-chr and -keep-allele-order.

#### qpGraph

qpGraph is part of ADMIXTOOLS software ^84^ and we used it to reconstruct the different relationships across the species in the study by comparing the various f statistics (f2, f3, f4) and generating an admixture graph with the best fitting admixture proportions and branch length (in unit of genetic drift). In particular, the graph generated using this tool is the representation of the relationship between 6 main lineages: Sardinian dhole, Asian dhole, AHDs, Ethiopian wolf and grey wolves.

In order to run qpGraph the PLINK files (BED BIM and FAM) were converted into EIGENSTRAT format using the package ConvertF implemented in ADMIXTOOLS. Then a qpGraph par file was created to specify working directory, the genotypename, snpname and indivname files. The options hires: YES, lsqmode: YES, blgsize: 0.005 and diag: .0001 were applied.

The samples were clustered into different groups representing the main lineages: AHDs, Asian dholes, Sardinian dhole, Ethiopian wolf and grey wolf. The best strategy was to start with a small graph and later add the other populations. Therefore, we built a graph topology including only the outgroup (the Andean fox), 6 AHDs, 2 Asian dholes, and the Sardinian dhole. When constructing the graph, it is important to do so considering the tree topology already available (see ASTRAL and DiscoVista trees) in order to specify the *root*, the outgroup and every other lineage representing a leaf of the graph and connected through nodes. A graph with a Z-value within 3 and −3 was considered significant, meaning that the topology of the graph fits with the combination of the f-statistics. Instead, when the Z-value is higher than 3 or lower than −3 the topology does not fully represent the relationship between the samples in the study and changes are necessary. Therefore when constructing the graph, the Sardinian dhole was modelled as sister clade to all possible internal and external nodes and as admixed from different node pairs and the Z values were then evaluated. Subsequently, once confirmed that the graph had a good fit (− 3 > Z < +3) we added the Ethiopian wolf and later a grey wolf (Portuguese wolf) and evaluated their fit in the phylogeny as explained above.

#### Split time analysis - F(A|B)

We also investigated the divergence time between the two dhole lineages by computing the probability F(A|B) that an (ancient) individual A (in this case the Sardinian dhole) carries a derived heterozygous allele in an individual B (Asian dhole). We estimated the standard error using a block jackknife estimate of the statistic, using a block size of 1 MB to partition the genome into non-overlapping regions. The assumption behind this approach is that when two populations start to diverge, they will also accumulate mutations that - due to isolation - will not be shared with other populations. Therefore, we first polarized the alleles to ancestral and derived using the Andean fox genome and we then called haplotypes with minimum base quality of 25 for the population B that we used to select only the heterozygous sites. We used these sites to compute the probability that the individual A would carry the derived allele, by randomly sampling each allele. To calibrate the probability F(A|B) to the population size history of the Asian dhole, we computed PSMC on the higher coverage genome Asian dhole in our dataset (Beijing Zoo dhole). The PSMC inference was based on the parameters, -N25 - t15 -r5 -p “4+25*2+4+6”, and the output was used to simulate 900 mb through msHOT ^85,86^. Different divergence times were computed, every 10 ka, spanning from 30 ka to 400 ka (expressed in generation time). We found the divergence time range by identifying the intersection between the empirical value line and the expected decay of F(A|B) as a function of split time.

#### hPSMC

To estimate the end of gene flow between the Sardinian dhole and the Asian dhole we used hPSMC ^32^. We first used ANGSD to generate haploid consensus sequences mapped to the dog reference genome and considering only autosomes. Bases with base quality lower than 30 and reads with mapping quality lower than 30 were discarded (-minQ 30 -minmapq 30). The minimum depth was set to 2x (-setminDepth 2) and the following quality filters: - remove_bads 1, -uniqueOnly 1, -baq 1 and -C 50. The two fasta files generated were combined into a diploid sequence using the hPSMC tool psmcfa_from_2_fastas.py. Subsequently, we ran the psmcfa output through PSMC with the parameters (-p) “4+25*2+4+6”, number of iterations = 25 (-N25), maximum 2N0 coalescent time = 15 (-t15), initial theta/rho ratio = 5 (-r5). We used psmc_plot.pl to translate into a plot this information and we assumed a mutation rate of 4 × 10^−9^ per base pair per generation ^30^ and 3 years generation time ^30,31^. The pre-divergence effective population size (Ne) estimated from the output was used to run simulations using hPSMC_quantify_split_time.py script from hPSMC tool with different divergence times, between 100,000 and 700 ka in 50 ka intervals using ms ^85^.

## Acknowledgments

This research was funded by the ERC Consolidator Grant 681396 ‘Extinction Genomics’. This work has been performed within a five-year (2017-2022) scientific agreement between the “Soprintendenza Archeologia, Belle arti e Paesaggio per le province di Sassari, Olbia-Tempio e Nuoro” and the University of Florence Earth Sciences Dept., and is framed within a wider project on Late Neogene vertebrate evolution granted by the University of Florence (Fondi di Ateneo) under the responsibility of L.R..

The authors also acknowledge support from Science for Life Laboratory, the Knut and Alice Wallenberg Foundation, the National Genomics Infrastructure funded by the Swedish Research Council, and Uppsala Multidisciplinary Center for Advanced Computational Science for assistance with massively parallel sequencing and access to the UPPMAX computational infrastructure. We would like to thank Davide Palumbo, Elisabetta Cilli and Romolo Caniglia for useful intellectual discussions about this research.

## Author Contributions

Conceptualization, M.M.C., M.H.S.S., S.G. and M.T.P.G.;

Investigation, M.M.C., S.G.;

Formal Analysis, M.M.C.; J.K.J.;

Data Curation, M.M.C., J.K.J.;

Resources, C.T., E.C., E.R., L.D., L.R., G.Z., C.G., M.A., M.T.P.G., O.S., T.S.P., S.B.L. and S.G.;

Writing, Original Draft, M.M.C.;

Writing -Review & Editing, M.M.C., M.H.S.S., S.B.L., E.R., L.R., C.T., M.A., E.C., M.G., Et.R., L.D., M.T.P.G. and S.G.;

Visualization, M.M.C.;

Supervision, S.G.;

Funding Acquisition, M.T.P.G

## Resource Availability

### Lead contact

Further information and requests for reagents and data may be directed to and will be fulfilled by the Lead Contact, Shyam Gopalakrishnan

### Materials Availability

This study did not generate new unique reagents.

### Data and Code Availability

Raw fastq reads for nuclear data of the Sardinian dhole will be deposited at the European Nucleotide Archive (ENA; study accession number ——) before publication.

## References

1. Malatesta, A. (1970). Cynotherium sardous Studiati an extinct canid from the Pleistocene of Sardinia. Mem. dell’Istituti Ital. di Paleontol. Umana, NS 1, 1–72.

2. Vigne, J.-D., Bailon, S., and Cuisin, J. (1997). Biostratigraphy of Amphibians, Reptiles, Birds and Mammals in Corsica and the role of Man in the Holocene faunal turnover. Anthropozoologica 25-26SP-587, 604.

3. Lyras, G.A., van der Geer, A.A.E., Dermitzakis, M.D., and De Vos, J. (2006). Cynotherium sardous, an insular canid (Mammalia: Carnivora) from the Pleistocene of Sardinia (Italy), and its origin. Journal of Vertebrate Paleontology 26, 735–745.

4. Palombo, M.R. (2018). Insular mammalian fauna dynamics and paleogeography: A lesson from the Western Mediterranean islands. Integr. Zool. 13, 2–20.

5. Palombo, M.R. (2006). Biochronology of the Plio-Pleistocene terrestrial mammals of Sardinia: the state of the art. Hellenic Journal of Geosciences 41, 47–66.

6. Eisenmann, V., and van der Geer, B. (1999). The Cynotherium from Corbeddu (Sardinia): comparative biometry with extant and fossil canids. Deinsea 7, 147–168.

7. Abbazzi, L., Angelone, C., Arca, M., Barisone, G., Bedetti, C., Delfino, M., Kotsakis, T., Marcolini, F., Palombo, M.R., Pavia, M., et al. (2004). Plio-Pleistocene fossil vertebrates of Monte Tuttavista (Orosei, Eastern Sardinia, Italy), an overview. Rivista Italiana di Paleontologia e Stratigrafia 110, 681–706.

8. Lyras, G., and van der Geer, A. (2006). Adaptations of the Pleistocene island canid Cynotherium sardous (Sardinia, Italy) for hunting small prey. Cranium 23, 51–60.

9. Madurell-Malapeira, J., Palombo, M.R., and Sotnikova, M. (2015). Cynotherium malatestai, sp. Nov. (Carnivora, Canidae) from the early middle Pleistocene deposits of Grotta dei Fiori (Sardinia, Western Mediterranean). J. Vert. Paleontol. 35.

10. Malatesta, A. (1962). Il cane selvaggio del Pleistocene di Sardegna.

11. Bonifay, M.-F. (1971). Carnivores quaternaires du Sud-Est de la France.

12. Abbazzi, L., Arca, M., Tuveri, C., and Rook, L. (2005). The endemic canid Cynotherium (Mammalia, Carnivora) from the Pleistocene deposits of Monte Tuttavista (Nuoro, Eastern Sardinia). Rivista Italiana di Paleontologia e Stratigrafia 111, 497–511.

13. Gopalakrishnan, S., Sinding, M.-H.S., Ramos-Madrigal, J., Niemann, J., Samaniego Castruita, J.A., Vieira, F.G., Carøe, C., de Manuel Montero, M., Kuderna, L., Serres, A., et al. (2019). Interspecific Gene Flow Shaped the Evolution of the Genus Canis. Curr. Biol. 29, 4152.

14. Zhang, C., Rabiee, M., Sayyari, E., and Mirarab, S. (2018). ASTRAL-III: polynomial time species tree reconstruction from partially resolved gene trees. BMC Bioinformatics 19, 153.

15. Allman, E.S., Degnan, J.H., and Rhodes, J.A. (2011). Identifying the rooted species tree from the distribution of unrooted gene trees under the coalescent. J. Math. Biol. 62, 833–862.

16. Krijgsman, W., Hilgen, F.J., Raffi, I., Sierro, F.J., and Wilson, D.S. (1999). Chronology, causes and progression of the Messinian salinity crisis. Nature 400, 652–655.

17. Palombo, M.R., and Rozzi, R. (2014). How correct is any chronological ordering of the Quaternary Sardinian mammalian assemblages? Quat. Int. 328-329, 136–155.

18. Sondaar, P.Y., Sanges, M., Kotsakis, T., and de Boer, P.L. (1986). The Pleistocene deer hunter of Sardinia. Geobios Mem. Spec. 19, 17–31.

19. Palombo, M.R. (2009). Biochronology, paleobiogeography and faunal turnover in western Mediterranean Cenozoic mammals. Integr. Zool. 4, 367–386.

20. Melis, R.T., Palombo, M.R., Ghaleb, B., and Meloni, S. (2016). A key site for inferring the timing of dispersal of giant deer in Sardinia, the Su Fossu de Cannas cave, Sadali, Italy. Quat. Res. 86, 335–347.

21. Roy, M.S., Girman, D.J., Taylor, A.C., and Wayne, R.K. (1994). The use of museum specimens to reconstruct the genetic variability and relationships of extinct populations. Experientia 50, 551–557.

22. Gottelli, D., Sillero-Zubiri, C., Marino, J., Funk, S.M., and Wang, J. (2013). Genetic structure and patterns of gene flow among populations of the endangered Ethiopian wolf. Anim. Conserv. 16, 234–247.

23. Woodroffe, R., and Ginsberg, J.R. (1999). Conserving the African wild dog Lycaon pictus. II. Is there a role for reintroduction? Oryx 33, 143–151.

24. Marsden, C.D., Woodroffe, R., Mills, M.G.L., McNutt, J.W., Creel, S., Groom, R., Emmanuel, M., Cleaveland, S., Kat, P., Rasmussen, G.S.A., et al. (2012). Spatial and temporal patterns of neutral and adaptive genetic variation in the endangered African wild dog (Lycaon pictus): SPATIAL AND TEMPORAL DIVERSITY IN WILD DOGS. Mol. Ecol. 21, 1379–1393.

25. Zimen, E., and Boitani, L. (1975). Number and distribution of wolves in Italy. Z. Säugetierkunde 40, 102–112.

26. Kamler, J., Institution), N.S. (smithsonian, Jenks, K., Srivathsa, A., Sheng, L., and Kunkel, K. (2015). IUCN Red List of Threatened Species: Cuon alpinus. IUCN Red List of Threatened Species.

27. Campana, M.G., Parker, L.D., Hawkins, M.T.R., Young, H.S., Helgen, K.M., Szykman Gunther, M., Woodroffe, R., Maldonado, J.E., and Fleischer, R.C. (2016). Genome sequence, population history, and pelage genetics of the endangered African wild dog (Lycaon pictus). BMC Genomics 17, 1013.

28. Armstrong, E.E., Taylor, R.W., Prost, S., Blinston, P., van der Meer, E., Madzikanda, H., Mufute, O., Mandisodza-Chikerema, R., Stuelpnagel, J., Sillero-Zubiri, C., et al. (2019). Cost-effective assembly of the African wild dog (Lycaon pictus) genome using linked reads. Gigascience 8.

29. van der Valk, T., Díez-del-Molino, D., Marques-Bonet, T., Guschanski, K., and Dalén, L. (2019). Historical Genomes Reveal the Genomic Consequences of Recent Population Decline in Eastern Gorillas. Curr. Biol. 29, 165–170.e6.

30. Skoglund, P., Ersmark, E., Palkopoulou, E., and Dalén, L. (2015). Ancient wolf genome reveals an early divergence of domestic dog ancestors and admixture into high-latitude breeds. Curr. Biol. 25, 1515–1519.

31. Freedman, A.H., Gronau, I., Schweizer, R.M., Ortega-Del Vecchyo, D., Han, E., Silva, P.M., Galaverni, M., Fan, Z., Marx, P., Lorente-Galdos, B., et al. (2014). Genome Sequencing Highlights the Dynamic Early History of Dogs. PLoS Genet. 10, e1004016.

32. Cahill, J.A., Soares, A.E.R., Green, R.E., and Shapiro, B. (2016). Inferring species divergence times using pairwise sequential markovian coalescent modelling and low-coverage genomic data. Philos. Trans. R. Soc. Lond. B Biol. Sci. 371.

33. Van der Made, J. (1999). Biogeography and stratigraphy of the Mio-Pleistocene mammals of Sardinia and the description of some fossils. Deinsea 7, 337–360.

34. Madurell-Malapeira, J., Rook, L., Martínez-Navarro, B., Alba, D.M., Aurell-Garrido, J., and Moyà-solà, S. (2013). The latest European painted dog. J. Vert. Paleontol. 33, 1244–1249.

35. Stiner, M.C., Howell, F.C., Martínez-Navarro, B., Tchernov, E., and Bar-Yosef, O. (2001). Outside Africa: Middle Pleistocene Lycaon from Hayonim Cave, Israel. BOLLETTINO-SOCIETA PALEONTOLOGICA ITALIANA 40, 293–302.

36. Schule, W. (1993). Mammals, Vegetation and the Initial Human Settlement of the Mediterranean Islands: A Palaeoecological Approach. J. Biogeogr. 20, 399–411.

37. Gippoliti, S., and Amori, G. (2006). Ancient introductions of mammals in the Mediterranean Basin and their implications for conservation. Mamm. Rev. 36, 37–48.

38. Martini, F. (1992). Il più antico popolamento umano delle isole: la Sardegna. I primi abitanti della Valle Padana: Monte Poggiolo, Ed. Jaca Book, Milan, Italie, 175–187.

39. [No title] https://www.researchgate.net/profile/Rengert_Elburg/publication/215481837_The_human_colonization_of_Sardinia_a_Late-Pleistocene_human_fossil_from_Corbeddu_cave/links/09e4150aad8b1230c6000000.pdf.

40. Spoor, F. (1999). The human fossils from Corbeddu Cave, Sardinia: a reappraisal. Deinsea 7, 297–302.

41. Palombo, M.R., Antonioli, F., Presti, V.L., Mannino, M.A., Melis, R.T., Orru, P., Stocchi, P., Talamo, S., Quarta, G., Calcagnile, L., et al. (2017). The late Pleistocene to Holocene palaeogeographic evolution of the Porto Conte area: Clues for a better understanding of human colonization of Sardinia and faunal dynamics during the last 30 ka. Quat. Int. 439, 117–140.

42. Hofmeijer, G.K. (1997). Late Pleistocene Deer Fossils from Corbeddu cave: implications for human Colonization of the Island of Sardinia (British Archaeological Reports Ltd).

43. Boessenkool, S., Hanghøj, K., Nistelberger, H.M., Der Sarkissian, C., Gondek, A.T., Orlando, L., Barrett, J.H., and Star, B. (2017). Combining bleach and mild predigestion improves ancient DNA recovery from bones. Mol. Ecol. Resour. 17, 742–751.

44. Allentoft, M.E., Sikora, M., Sjögren, K.-G., Rasmussen, S., Rasmussen, M., Stenderup, J., Damgaard, P.B., Schroeder, H., Ahlström, T., Vinner, L., et al. (2015). Population genomics of Bronze Age Eurasia. Nature 522, 167–172.

45. Carøe, C., Gopalakrishnan, S., Vinner, L., Mak, S.S.T., Sinding, M.H.S., Samaniego, J.A., Wales, N., Sicheritz-Pontén, T., and Gilbert, M.T.P. (2018). Single-tube library preparation for degraded DNA. Methods Ecol. Evol. 9, 410–419.

46. Mak, S.S.T., Gopalakrishnan, S., Carøe, C., Geng, C., Liu, S., Sinding, M.-H.S., Kuderna, L.F.K., Zhang, W., Fu, S., Vieira, F.G., et al. (2017). Comparative performance of the BGISEQ-500 vs Illumina HiSeq2500 sequencing platforms for palaeogenomic sequencing. Gigascience 6, 1–13.

47. Reimer, P.J., Austin, W.E.N., Bard, E., Bayliss, A., Blackwell, P.G., Ramsey, C.B., Butzin, M., Cheng, H.,Lawrence Edwards, R., Friedrich, M., et al. (2020). The IntCal20 Northern Hemisphere Radiocarbon Age Calibration Curve (0-55 cal kBP). Radiocarbon 62, 725–757.

48. Ramsey, C.B. (2009). Bayesian Analysis of Radiocarbon Dates. Radiocarbon 51, 337–360.

49. Auton, A., Rui Li, Y., Kidd, J., Oliveira, K., Nadel, J., Holloway, J.K., Hayward, J.J., Cohen, P.E., Greally, J.M., Wang, J., et al. (2013). Genetic recombination is targeted towards gene promoter regions in dogs. PLoS Genet. 9, e1003984.

50. Wang, G.-D., Zhai, W., Yang, H.-C., Fan, R.-X., Cao, X., Zhong, L., Wang, L., Liu, F., Wu, H., Cheng, L.-G., et al. (2013). The genomics of selection in dogs and the parallel evolution between dogs and humans. Nat. Commun. 4, 1860.

51. Zhang, W., Fan, Z., Han, E., Hou, R., Zhang, L., Galaverni, M., Huang, J., Liu, H., Silva, P., Li, P., et al.(2014). Hypoxia adaptations in the grey wolf (Canis lupus chanco) from Qinghai-Tibet Plateau. PLoS Genet. 10, e1004466.

52. Wang, G.-D., Zhai, W., Yang, H.-C., Wang, L., Zhong, L., Liu, Y.-H., Fan, R.-X., Yin, T.-T., Zhu, C.-L., Poyarkov, A.D., et al. (2016). Out of southern East Asia: the natural history of domestic dogs across the world. Cell Res. 26, 21–33.

53. Liu, Y.-H., Wang, L., Xu, T., Guo, X., Li, Y., Yin, T.-T., Yang, H.-C., Hu, Y., Adeola, A.C., Sanke, O.J., et al. (2018). Whole-Genome Sequencing of African Dogs Provides Insights into Adaptations against Tropical Parasites. Mol. Biol. Evol. 35, 287–298.

54. Koepfli, K.-P., Pollinger, J., Godinho, R., Robinson, J., Lea, A., Hendricks, S., Schweizer, R.M., Thalmann, O., Silva, P., Fan, Z., et al. (2015). Genome-wide Evidence Reveals that African and Eurasian Golden Jackals Are Distinct Species. Curr. Biol. 25, 2158–2165.

55. Fan, Z., Silva, P., Gronau, I., Wang, S., Armero, A.S., Schweizer, R.M., Ramirez, O., Pollinger, J., Galaverni, M., Del-Vecchyo, D.O., et al. (2016). Worldwide patterns of genomic variation and admixture in gray wolves. Genome Res. 26, 163–173.

56. Sinding, M.-H.S., Gopalakrishan, S., Vieira, F.G., Samaniego Castruita, J.A., Raundrup, K., Heide Jørgensen, M.P., Meldgaard, M., Petersen, B., Sicheritz-Ponten, T., Mikkelsen, J.B., et al. (2018). Population genomics of grey wolves and wolf-like canids in North America. PLoS Genet. 14, e1007745.

57. vonHoldt, B.M., Cahill, J.A., Fan, Z., Gronau, I., Robinson, J., Pollinger, J.P., Shapiro, B., Wall, J., and Wayne, R.K. (2016). Whole-genome sequence analysis shows that two endemic species of North American wolf are admixtures of the coyote and gray wolf. Sci Adv 2, e1501714.

58. Schubert, M., Ermini, L., Der Sarkissian, C., Jónsson, H., Ginolhac, A., Schaefer, R., Martin, M.D., Fernández, R., Kircher, M., McCue, M., et al. (2014). Characterization of ancient and modern genomes by SNP detection and phylogenomic and metagenomic analysis using PALEOMIX. Nat. Protoc. 9, 1056–1082.

59. Schubert, M., Lindgreen, S., and Orlando, L. (2016). AdapterRemoval v2: rapid adapter trimming, identification, and read merging. BMC Res. Notes 9, 88.

60. Gopalakrishnan, S., Samaniego Castruita, J.A., Sinding, M.-H.S., Kuderna, L.F.K., Räikkönen, J., Petersen, B., Sicheritz-Ponten, T., Larson, G., Orlando, L., Marques-Bonet, T., et al. (2017). The wolf reference genome sequence (Canis lupus lupus) and its implications for Canis spp. population genomics. BMC Genomics 18, 495.

61. Lindblad-Toh, K., Wade, C.M., Mikkelsen, T.S., Karlsson, E.K., Jaffe, D.B., Kamal, M., Clamp, M., Chang, J.L., Kulbokas, E.J., 3rd, Zody, M.C., et al. (2005). Genome sequence, comparative analysis and haplotype structure of the domestic dog. Nature 438, 803–819.

62. Li, H., and Durbin, R. (2009). Fast and accurate short read alignment with Burrows-Wheeler transform. Bioinformatics 25, 1754–1760.

63. Toolkit, P. (2018). Picard Toolkit. Broad Institute.

64. DePristo, M.A., Banks, E., Poplin, R., Garimella, K.V., Maguire, J.R., Hartl, C., Philippakis, A.A., del Angel, G., Rivas, M.A., Hanna, M., et al. (2011). A framework for variation discovery and genotyping using next-generation DNA sequencing data. Nat. Genet. 43, 491–498.

65. McKenna, A., Hanna, M., Banks, E., Sivachenko, A., Cibulskis, K., Kernytsky, A., Garimella, K., Altshuler, D., Gabriel, S., Daly, M., et al. (2010). The Genome Analysis Toolkit: a MapReduce framework for analyzing next-generation DNA sequencing data. Genome Res. 20, 1297–1303.

66. Jónsson, H., Ginolhac, A., Schubert, M., Johnson, P.L.F., and Orlando, L. (2013). mapDamage2.0: fast approximate Bayesian estimates of ancient DNA damage parameters. Bioinformatics 29, 1682–1684.

67. Allaire, J. (2012). RStudio: integrated development environment for R. Boston, MA.

68. RStudio Team (2020). RStudio: Integrated Development Environment for R.

69. Wickham, H. (2016). ggplot2: Elegant Graphics for Data Analysis (Springer-Verlag New York).

70. Korneliussen, T.S., Albrechtsen, A., and Nielsen, R. (2014). ANGSD: Analysis of Next Generation Sequencing Data. BMC Bioinformatics 15, 356.

71. Meisner, J., and Albrechtsen, A. (2018). Inferring Population Structure and Admixture Proportions in Low-Depth NGS Data. Genetics 210, 719–731.

72. Skotte, L., Korneliussen, T.S., and Albrechtsen, A. (2013). Estimating individual admixture proportions from next generation sequencing data. Genetics 195, 693–702.

73. Behr, A.A., Liu, K.Z., Liu-Fang, G., Nakka, P., and Ramachandran, S. (2016). pong: fast analysis and visualization of latent clusters in population genetic data. Bioinformatics 32, 2817–2823.

74. Hadley Wickham, R.F., Henry, L., Müller, K., and Others (2018). dplyr: A grammar of data manipulation. Version 0. 7 6.

75. Quinlan, A.R., and Hall, I.M. (2010). BEDTools: a flexible suite of utilities for comparing genomic features. Bioinformatics 26, 841–842.

76. Li, H., Handsaker, B., Wysoker, A., Fennell, T., Ruan, J., Homer, N., Marth, G., Abecasis, G., Durbin, R., and 1000 Genome Project Data Processing Subgroup (2009). The Sequence Alignment/Map format and SAMtools. Bioinformatics 25, 2078–2079.

77. Kozlov, A.M., Darriba, D., Flouri, T., Morel, B., and Stamatakis, A. (2019). RAxML-NG: a fast, scalable and user-friendly tool for maximum likelihood phylogenetic inference. Bioinformatics 35, 4453–4455.

78. Letunic, I., and Bork, P. (2019). Interactive Tree Of Life (iTOL) v4: recent updates and new developments. Nucleic Acids Res. 47, W256–W259.

79. Sayyari, E., Whitfield, J.B., and Mirarab, S. (2018). DiscoVista: Interpretable visualizations of gene tree discordance. Mol. Phylogenet. Evol. 122, 110–115.

80. Poplin, R., Ruano-Rubio, V., DePristo, M., Fennell, T., Carneiro, M., Van der Auwera, G., Kling, D., Gauthier, L., Levy-Moonshine, A., Roazen, D., et al. (2017). Scaling accurate genetic variant discovery to tens of thousands of samples. bioRxiv, 201178.

81. Danecek, P., Auton, A., Abecasis, G., Albers, C.A., Banks, E., DePristo, M.A., Handsaker, R.E., Lunter, G., Marth, G.T., Sherry, S.T., et al. (2011). The variant call format and VCFtools. Bioinformatics 27, 2156–2158.

82. Purcell, S., Neale, B., Todd-Brown, K., Thomas, L., Ferreira, M.A.R., Bender, D., Maller, J., Sklar, P., deBakker, P.I.W., Daly, M.J., et al. (2007). PLINK: a tool set for whole-genome association and population based linkage analyses. Am. J. Hum. Genet. 81, 559–575.

83. Chang, C.C., Chow, C.C., Tellier, L.C.A.M., Vattikuti, S., Purcell, S.M., and Lee, J.J. (2015). Second-generation PLINK: rising to the challenge of larger and richer datasets. Gigascience 4.

84. Patterson, N., Moorjani, P., Luo, Y., Mallick, S., Rohland, N., Zhan, Y., Genschoreck, T., Webster, T., and Reich, D. (2012). Ancient admixture in human history. Genetics 192, 1065–1093.

85. Hudson, R.R. (2002). Generating samples under a Wright-Fisher neutral model of genetic variation. Bioinformatics 18, 337–338.

86. Hellenthal, G., and Stephens, M. (2007). msHOT: modifying Hudson’s ms simulator to incorporate crossover and gene conversion hotspots. Bioinformatics 23, 520–521.

