## Supplemental Data for "Genomic analyses of the extinct Sardinian dhole (*Cynotherium sardous*) reveal its evolutionary history"

**Authors affiliation:**

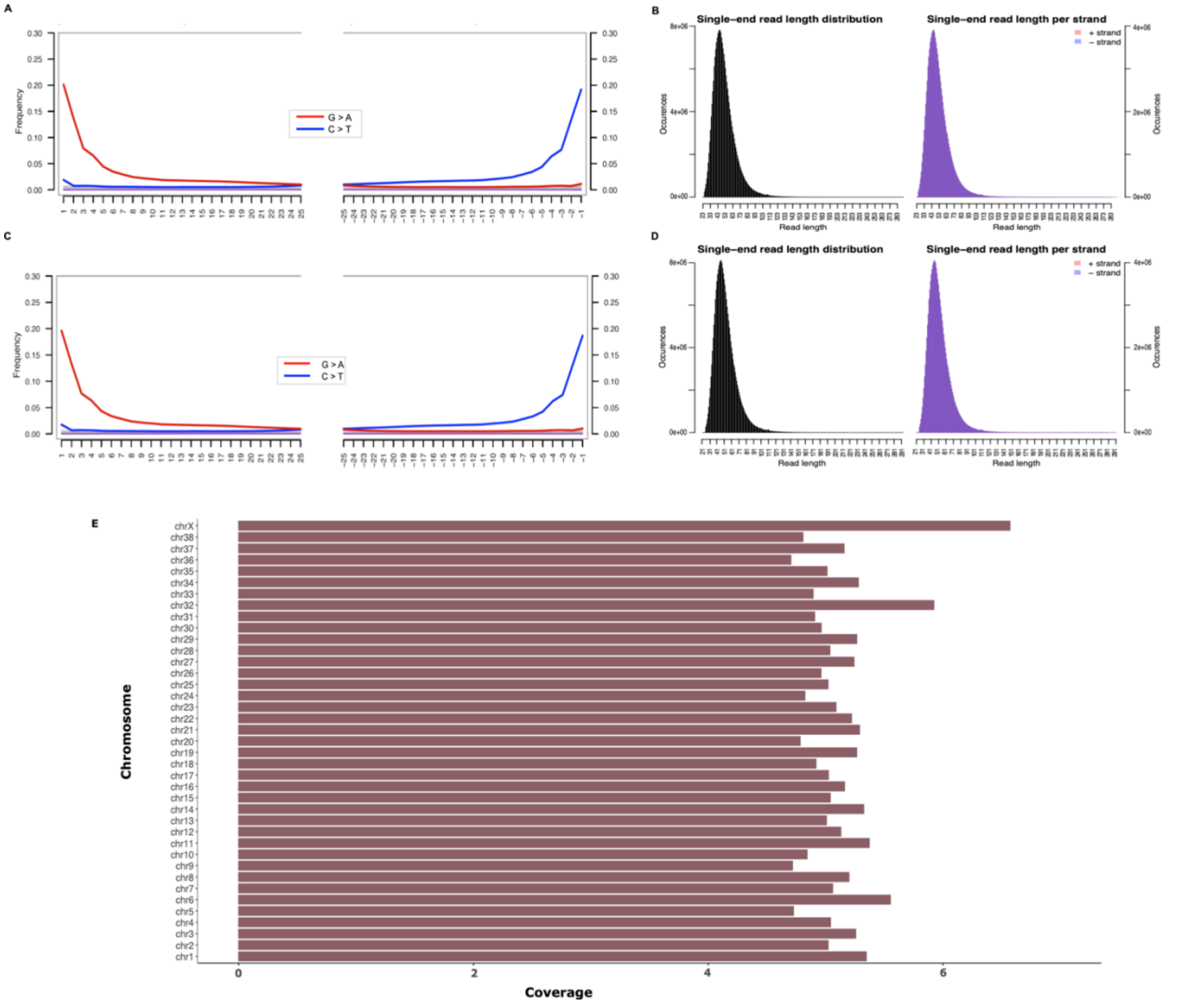


**Figure S1. Nucleotide misincorporation rates, length distribution and chromosome coverage of the Sardinian dhole.**

A) Nucleotide misincorporation rates of the sample mapped to the wolf reference genome.
B) Length distribution of the total reads mapped to the wolf reference genome.
C) Nucleotide misincorporation rates of the sample mapped to the CanFam3.1 reference genome.
D) Length distribution of the total reads mapped to the CanFam3.1 reference genome.
E) Biological sex determination. The sequencing reads sample were mapped to the dog reference genome (CanFam3.1)[1], which present the genome divided in 38 autosomal chromosomes and the sex chromosome X. The plot here shows on the x axis the average depth of coverage at each chromosome of the Sardinian dhole which resulted to be ~ 5x across the whole genome. However, the average depth of coverage for the chromosome X was higher (> 6x) than the average depth of each chromosome and the whole genome, thus we concluded that our sample was female.

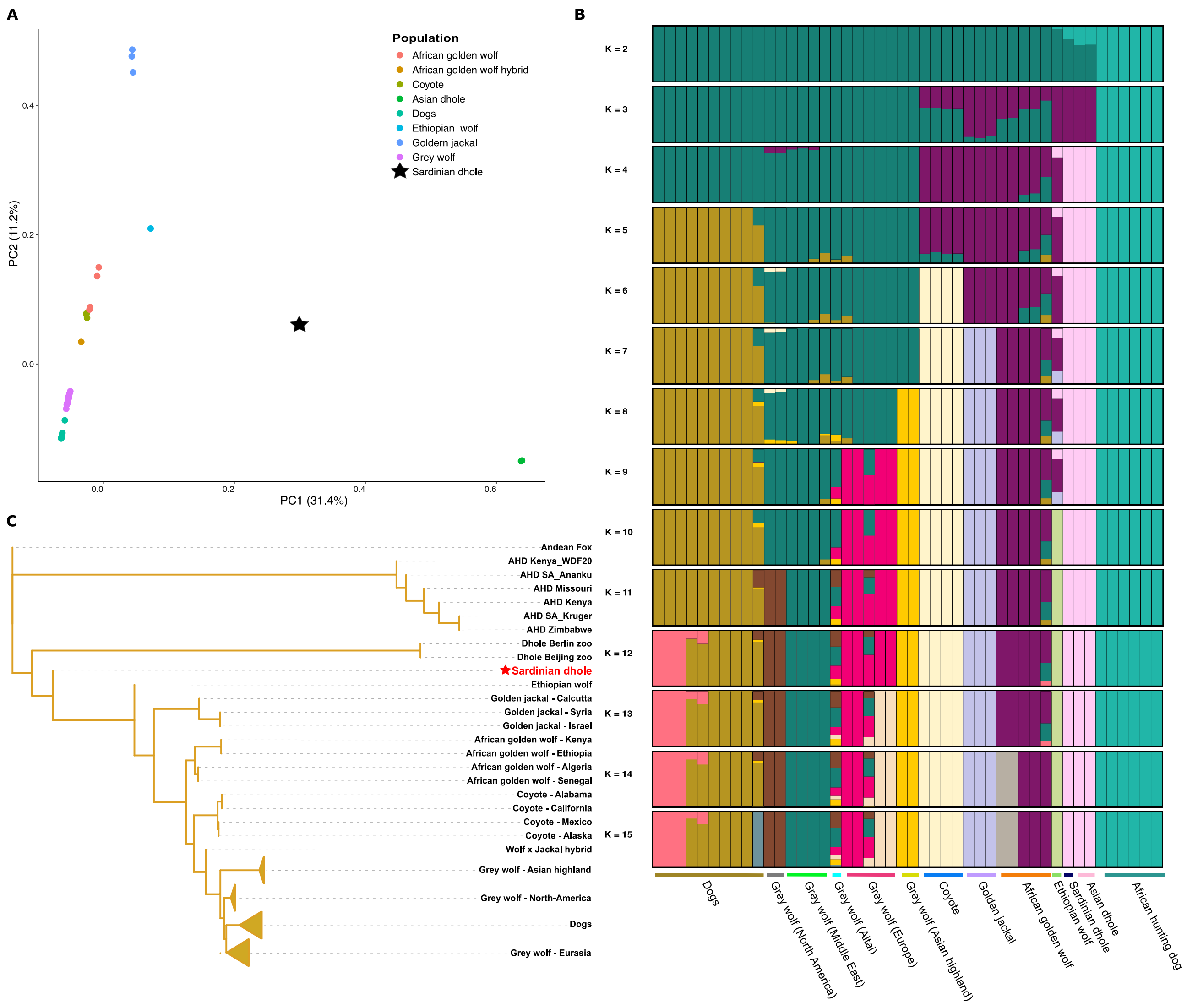


**Figure S2. PCA, Admixture proportions, and Astral tree**

A) Principal component analysis (PCA) of 41 canids excluding Andean fox and the African hunting dogs

B) The admixture proportions were estimated using NGSAdmix[2], for a range of ancestry components from K=2 to K=15.

C) The phylogeny of 47 samples was computed by Astral III[3] on 1000 gene trees generated with RAxML-ng[4] under the evolutionary model GTR-GAMMA.

Dogs, Eurasian, North-American and Chinese highland wolf nodes were collapsed separately.

The node labels represent the posterior probability computed by RAxML-ng using 100 replicated in Astral-III. Branch nodes with a posterior probability above 0.75 are shown.

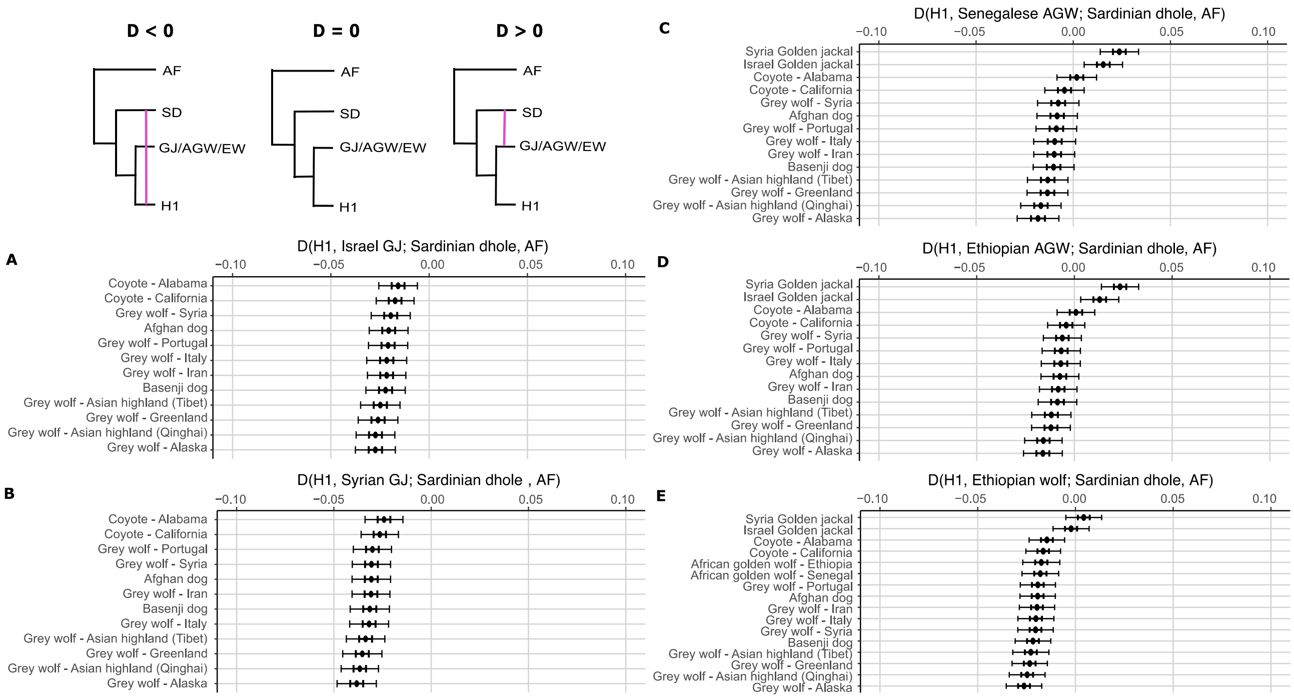


### Figure S3. D-stats

In this figure the gene flow among different canids is shown using ABBA-BABA test in ANGSD[5]. The following three different combinations were tested: 1) (((*Canis lupus*, Golden Jackal), Sardinian dhole), Andean fox), 2) (((*Canis lupus*, African golden wolf), Sardinian dhole), Andean fox) and 3) (((*Canis lupus*, Ethiopian wolf), Sardinian dhole), Andean fox).

In certain places in the figure the species names are abbreviated with AF, SD, GJ, AGW and EW for Andean for, Sardinian dhole, Golden jackal, African golden wolf and Ethiopian wolf respectively.

In the figure A-B) the first combination is shown and there is significant allele sharing (D < 0) between the coyotes, wolves and dogs to/from Sardinian dhole (SD) when considering two Golden jackals (GJ) (Syria and Israel golden jackals) in H2.

In panel C-D) the combination 2) was tested and the allele sharing between the AGW and SD when placing the GJ in H1. However, when considering *Canis lupus* in H1 the D-stats shift to a negative result (D < 0) indicating gene flow into the Sardinian dhole.

In figure E) the combination 3) is shown and when considering all the *Canis* crown species but Golden jackal (GJ) the D-statistics show a negative D (D < 0 - BABA) meaning that there is allele sharing between these species and the Sardinian dhole (SD).

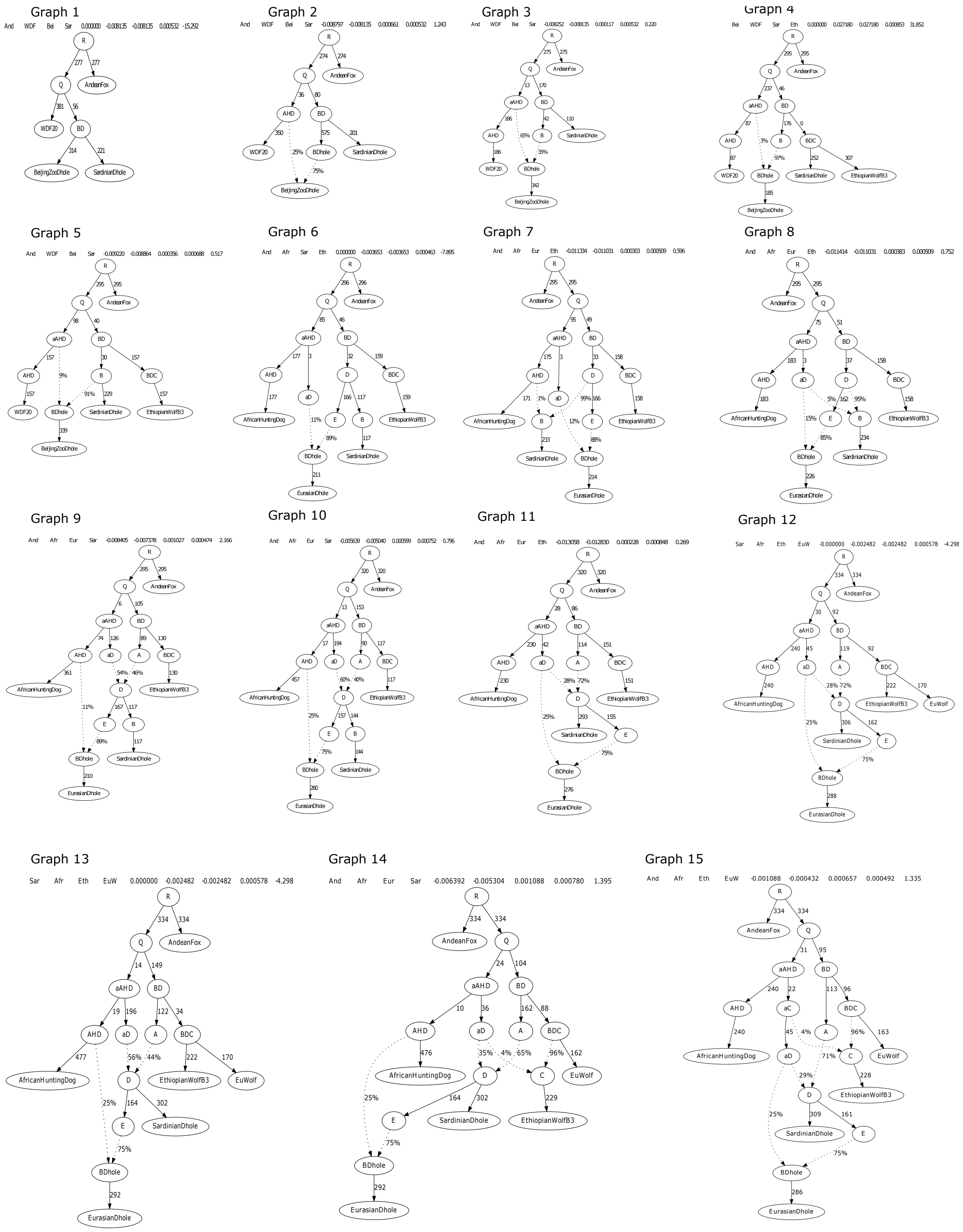


**Figure S4. Admixture graph population history models.** The demographic models of the Sardinian dhole, modern African hunting dogs, Eurasian dholes, Ethiopian wolves and Eurasian wolves, estimated using qpGraph of Admixtools[6] with the observed and expected f-statistics to compute the admixture graph**.**

Related to Figure 3E in the main article.

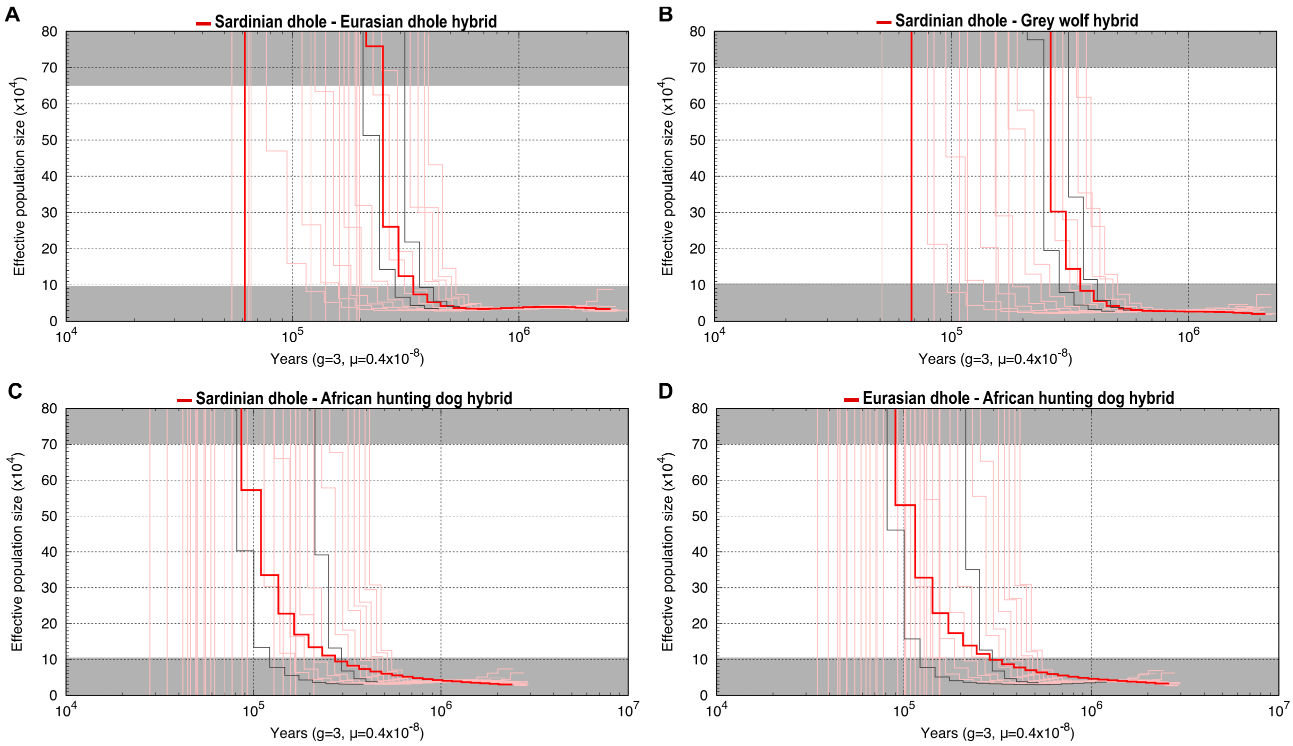


**Figure S5. Hybrid PSMC (hPSMC)[7] plot of the Sardinian and the Eurasian dholes with simulations of different divergence times spanning 100,000-700,000 years in 50,000 years intervals.**

Related to Figure 4 in the main text.

Greyed out regions represent 1.5x and 10x the pre-divergence effective population size. Red lines in bold represent the hPSMC results based on the real data while the remaining red thin lines represent the simulated data. The simulation closest to the real data that do not overlap with it are shown in dark grey and were used to infer the time range in which the gene flow ceased between A) the Sardinian dhole and the Eurasian dhole, B) the Sardinian dhole and European wolf (Portugal), C) the Sardinian dhole and African hunting dog (Zimbabwe) and D) the Eurasian dhole and African hunting dog (Zimbabwe).

### References


1. Lindblad-Toh, K., Wade, C.M., Mikkelsen, T.S., Karlsson, E.K., Jaffe, D.B., Kamal, M., Clamp, M., Chang, J.L., Kulbokas, E.J., 3rd, Zody, M.C., et al. (2005). Genome sequence, comparative analysis and haplotype structure of the domestic dog. Nature *438*, 803–819.

2. Skotte, L., Korneliussen, T.S., and Albrechtsen, A. (2013). Estimating individual admixture proportions from next generation sequencing data. Genetics *195*, 693–702.

3. Zhang, C., Rabiee, M., Sayyari, E., and Mirarab, S. (2018). ASTRAL-III: polynomial time species tree reconstruction from partially resolved gene trees. BMC Bioinformatics *19*, 153.

4. Kozlov, A.M., Darriba, D., Flouri, T., Morel, B., and Stamatakis, A. (2019). RAxML-NG: a fast, scalable and user-friendly tool for maximum likelihood phylogenetic inference. Bioinformatics *35*, 4453–4455.

5. Korneliussen, T.S., Albrechtsen, A., and Nielsen, R. (2014). ANGSD: Analysis of Next Generation Sequencing Data. BMC Bioinformatics *15*, 356.

6. Patterson, N., Moorjani, P., Luo, Y., Mallick, S., Rohland, N., Zhan, Y., Genschoreck, T., Webster, T., and Reich, D. (2012). Ancient admixture in human history. Genetics *192*, 1065–1093.

7. Cahill, J.A., Soares, A.E.R., Green, R.E., and Shapiro, B. (2016). Inferring species divergence times using pairwise sequential markovian coalescent modelling and low-coverage genomic data. Philos. Trans. R. Soc. Lond. B Biol. Sci. 371.
